# YeastMate: Neural network-assisted segmentation of mating and budding events in *S. cerevisiae*

**DOI:** 10.1101/2021.10.13.464238

**Authors:** David Bunk, Julian Moriasy, Felix Thoma, Christopher Jakubke, Christof Osman, David Hörl

## Abstract

Here, we introduce *YeastMate*, a user-friendly deep learning-based application for automated detection and segmentation of *Saccharomyces cerevisiae* cells and their mating and budding events in microscopy images. We build upon Mask R-CNN with a custom segmentation head for the subclassification of mother and daughter cells during lifecycle transitions. *YeastMate* can be used directly as a Python library or through a stand-alone GUI application and a Fiji plugin as easy to use frontends.

The source code for YeastMate is freely available at https://github.com/hoerlteam/YeastMate under the MIT license. We offer packaged installers for our whole software stack for Windows, macOS and Linux. A detailed user guide is available at https://yeastmate.readthedocs.io.

## Introduction

An important experimental approach when working with the budding yeast *Saccharomyces cerevisiae* is to examine the effects of mutations on morphological features or protein localization either by brightfield or fluorescent microscopy (1). Such experiments can also be performed in a systematic manner by combining automated microscopy with strain libraries comprising thousands of yeast strains that carry gene deletions or express fluorescently tagged proteins (2–4), which, however, calls for robust automated image analysis pipelines. In recent years, tools based on convolutional neural networks (CNNs) (5) have become state-of-the-art for many tasks in biomedical image anlysis (6), including segmentation of individual *S. cerevisiae* cells (7–9). Many experimental strategies facilitated by the yeast system also make use of specific transitions in the yeast lifecycle like budding and mating to study things like organelle inheritance and mitochondrial quality control (10, 11). Detection of matings and buddings is often done by hand or through dedicated postprocessing routines on the output of a single-cell segmentation tool (e.g., tracking of cells in a time series of images), highlighting the need for easy to use end-to-end solutions for these more complex tasks.

Here, we present *YeastMate*, a novel deep learning-based tool for end-to-end segmentation of single cells and detection of transitions in the lifecycle of *S. cerevisiae* in single transmitted light images. YeastMate performs three tasks: instance segmentation of single cells, object detection of zygotes and budding events, and automatic assignment of mother and daughter cells involved in a mating or budding event. The detection backend is based on Mask R-CNN (12) and is complemented by a user-friendly frontend using modern web technologies as well as a Fiji plugin. In the task of detecting mating and budding events, we achieve accuracies comparable to manual human re-annotation and we can also perform single cell segmentation robustly across various datasets. YeastMate is already being used in ongoing research in our lab (11). In addition to the software, we also provide a new dataset of images with manually annotated cells and mating and budding events.

## Materials and Methods

For a detailed description of our model architecture, postprocessing, software architecture and benchmarking strategies, as well as data generation and annotation, please refer to the Supplementary Methods.

### CNN architecture and training

Our network architecture builds upon Mask R-CNN with a modified mask segmentation head producing multiclass semenatic segmentations (Supplementary Figure S1) for each detected object. Instead of individually detecting mother and daughter cells, we first detect the whole mating and budding events as well as single cells. In a postprocessing step, we resolve the roles of the cells involved in budding or mating events based on the multiclass segmentation masks (Supplementary Figure S2). We used 80% of our images for training our network and cross-validation and 20% as a hold-out test set for the final performance assessment.

### Software architecture

YeastMate is implemented in a modular way: the detection backend can be used as a Python library but also runs as a webservice to provide its capabilities to client applications. As clients, we provide a Fiji plugin that directly interfaces with the server via HTTP requests as well as a standalone GUI Desktop application (Figure 1 and Supplementary Figures S3, S4, S5). We provide the whole YeastMate stack as a single installable file for use on a local workstation.

**Fig. 1.**
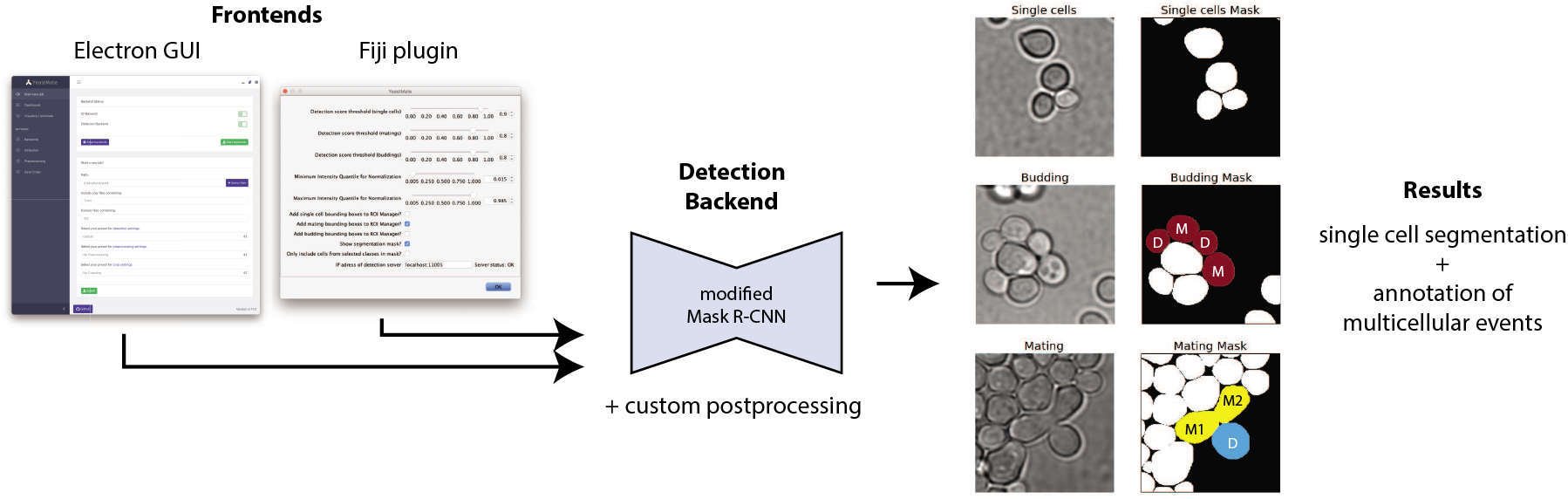
Main components and output of *YeastMate: We* perform instance segmentation of single cells and detection of lifecycle transitions using a modified Mask R-CNN (middle), which can either be used directly from Python code or via two GUI frontends (left) to provide instance segmentation of single cells as well as detection of budding events and mating events with identification of the mother (M) and daughter (D) cells involved in the event (right).

## Results

### Dataset

For training YeastMate, we collected 147 brightfield and differential interference contrast (DIC) images of *S. cerevisiae* acquired on two different microscopes at various imaging conditions and generated curated single cell segmentation masks and mating and budding annotations. In total, our data contain 17058 individual cells, 3615 buddings and 2380 zygotes (Supplementary Table ST1).

### Object detection performance

Our network achieves a mean average precision (mAP) of 0.878 as well as favourable APs for the individual classes of objects when applied to our test set (Supplementrary Figure S6). We also assessed inter-human reproducibility by comparing the annotations for mating and budding events of 4 different annotators (with the most experienced one chosen as the reference). On our test dataset, mean inter-human precision and recall are (0.908,0.946) for mating events and (0.774,0.763) for budding events. These values lie close to the PR-curves of our network, indicating that YeastMate can achieve object detection performance comparable to manual human annotation.

### Single cell segmentation performance

YeastMate also compares favourably to existing CNN-based solutions in single-cell segmentation, showing consistent performance not only on our own test dataset, but also two publicly available datasets (Supplementary Results, Supplementary Figure S7, Supplementary Table ST2).

## Conclusions

With YeastMate, we introduce an easy to use application to not only perform single cell segmentation in images of *S. cerevisiae* with high robustness across datasets, but also detect transitions in the cell cycle in single images with accuracies comparable to human annotation. YeastMate is implemented in a modular way and can be run on local workstations but the detection server can also be run on a remote compute server. Additionally, we provide two user-friendly frontends to make the tool available without the need to write code.

We can envision YeastMate to be expanded to detect other life cycle states, such as meiotic asci, similar to existing work in *A. thaliana* (13). YeastMate can provide a considerable improvement to high-throughput studies of yeast, facilitating not only the automated analysis of large image datasets of single cells, but also enabling the study the complex interplay of cellular components during both sexual and asexual reproduction of *S.cerevisiae*.

## ACKNOWLEDGEMENTS

The authors would like to thank Mikhail Slivinskiy (CHECK24 Vergleichsportal GmbH) for his helpful advice on the React.js frontend, as well as Heinrich Leonhardt and Hartmann Harz (LMU) for helpful comments on the manuscript and support. Some of the images in our dataset were acquired in the lab of Maya Schuldiner (Weizmann Institute) as part of an ongoing collaboration.

## Notes

https://github.com/hoerlteam/YeastMate

https://yeastmate.readthedocs.io/en/latest/

https://osf.io/287fr/?view_only=99d1fddb563b4253957f226c19c4113f

### Competing Interest Statement

The authors have declared no competing interest.

## Bibliography

1. Yoshikazu Ohya, Yoshitaka Kimori, Hiroki Okada, and Shinsuke Ohnuki. Single-cell phenomics in budding yeast. Molecular Biology of the Cell, 26(22):3920–3925, Nov 2015. ISSN 1059-1524, 1939-4586. doi: 10.1091/mbc.E15-07-0466.

2. Guri Giaever, Angela M. Chu, Li Ni, Carla Connelly, Linda Riles, Steeve Véronneau, Sally Dow, Ankuta Lucau-Danila, Keith Anderson, Bruno André, and et al. Functional profiling of the saccharomyces cerevisiae genome. Nature, 418(6896):387–391, Jul 2002. ISSN 0028-0836, 1476-4687. doi: 10.1038/nature00935.

3. Won-Ki Huh, James V Falvo, Luke C Gerke, Adam S Carroll, Russell W Howson, Jonathan S Weissman, and Erin K O’Shea. Global analysis of protein localization in budding yeast. Nature, 425(6959):686–691, 2003.

4. Uri Weill, Ido Yofe, Ehud Sass, Bram Stynen, Dan Davidi, Janani Natarajan, Reut Ben-Menachem, Zohar Avihou, Omer Goldman, Nofar Harpaz, et al. Genome-wide swap-tag yeast libraries for proteome exploration. Nature methods, 15(8):617–622, 2018.

5. Yann LeCun, Yoshua Bengio, and Geoffrey Hinton. Deep learning. nature, 521(7553): 436–444, 2015.

6. Lucas von Chamier, Romain F Laine, and Ricardo Henriques. Artificial intelligence for microscopy: what you should know. Biochemical Society Transactions, 47(4):1029–1040, 2019.

7. Alex X Lu, Taraneh Zarin, Ian S Hsu, and Alan M Moses. YeastSpotter: accurate and parameter-free web segmentation for microscopy images of yeast cells. Bioinformatics, 35 (21):4525–4527, May 2019. doi: 10.1093/bioinformatics/btz402.

8. Nicola Dietler, Matthias Minder, Vojislav Gligorovski, Augoustina Maria Economou, Denis Alain Henri Lucien Joly, Ahmad Sadeghi, Chun Hei Michael Chan, Mateusz Koziński, Martin Weigert, Anne-Florence Bitbol, et al. A convolutional neural network segments yeast microscopy images with high accuracy. Nature communications, 11 (1):1–8, 2020.

9. Danny Salem, Yifeng Li, Pengcheng Xi, Hilary Phenix, Miroslava Cuperlovic-Culf, and Mads Kaern. Yeastnet: Deep-learning-enabled accurate segmentation of budding yeast cells in bright-field microscopy. Applied Sciences, 11(6):2692, 2021.

10. Susanne M Rafelski, Matheus P Viana, Yi Zhang, Yee-Hung M Chan, Kurt S Thorn, Phoebe Yam, Jennifer C Fung, Hao Li, Luciano da F Costa, and Wallace F Marshall. Mitochondrial network size scaling in budding yeast. Science, 338(6108):822–824, 2012.

11. Christopher Jakubke, Rodaria Roussou, Andreas Maiser, Christina Schug, Felix Thoma, David Bunk, David Hörl, Heinrich Leonhardt, Peter Walter, Till Klecker, and Christof Osman. Cristae-dependent quality control of the mitochondrial genome. Science Advances, 7(36), 2021.

12. Kaiming He, Georgia Gkioxari, Piotr Dollar, and Ross Girshick. Mask r-cnn. In Proceedings of the IEEE International Conference on Computer Vision (ICCV), Oct 2017.

13. Eun-Cheon Lim, Jaeil Kim, Jihye Park, Eun-Jung Kim, Juhyun Kim, Yeong Mi Park, Hyun Seob Cho, Dohwan Byun, Ian R Henderson, Gregory P Copenhaver, et al. Deeptetrad: high-throughput image analysis of meiotic tetrads by deep learning in arabidopsis thaliana. The Plant Journal, 101(2):473–483, 2020.

