## Supplementary Material for "YeastMate: Neural network-assisted segmentation of mating and budding events in *S. cerevisiae*"

### Supplementary Methods

#### Data acquisition and cell culture

The images in our dataset are a collection of DIC and brightfield images from multiple acquisitions on either a Nikon Ti2 microscope equipped with a CFI SR HP Apo TIRF 100xH NA 1.49 oil immersion objective (with an optional 1.5x magnification relay lens) and dual Photometrics Prime 95B 25mm cameras or on a fully automated Olympus IX 83 microscope with an UPLFLM 60x NA 0.9 air objective with a Hamamatsu ORCA-Flash4.0 V3 camera. All together, we assembled our dataset from 6 individual experiments on the Nikon system and parts of 2 multi-well experiments imaged at the Olympus system. In all experiments, two haploid *S. cerevisiae* strains were mated. Data from the Nikon system were acquired for (Jakubke *et al.*, 2021), data from the Olympus system are preliminary data for a high-throughput screen. For a complete reference of the individual experiments included in the dataset, refer to Supplementary Table ST1.

Yeast cultivation and sample preparation was performed as described previously (Jakubke *et al.*, 2021). In short, to image the mating process, yeast cells of opposing mating type were grown in separate flasks in YPD medium at 30°C to an OD600 ~1.0. 1000 µl from both cultures were combined in a reaction tube, vortexed thoroughly and centrifuged for 3 minutes at 5.000 rpm. The pellet was resuspended in 50 µl of YPD medium and spotted onto a YPD plate to allow cells to mate. After incubation at 30°C for 3 h, pre-mated cells were scraped off and resuspended in 200µl PBS. The cell suspension was added into wells of 8-well IBIDI slides that had previously been coated with Concanavalin A. Attachment of cells to the coverslip was supported by centrifuging the IBIDI slide for 60 s at 1000 rpm.

#### Generation of ground truth annotations

Single-cell segmentation masks were generated in an iterative, feedback-based process: We initially trained a standard Mask R-CNN on the brightfield dataset provided by YeaZ (Dietler *et al.*, 2020), applied it to our images and manually corrected the generated masks using napari (Sofroniew *et al.*, 2021). For the annotation of cells involved in mating events or buddings, we implemented a custom annotation scheme, again using napari: For annotation of mating events, we manually drew paths in a shape layer going from mother 1 to mother 2 and, optionally, the medial daughter cell for each mating event in our images. Likewise, in another layer, we added paths going from mother to daughter cell in all buddings (Supplementary Figure S4, annotations are visualized in this manner in Supplementary Figure S6B). Our path-based annotations were saved in JSON files and were converted to the bounding box annotations needed by R-CNN-based networks on the fly when training the network.

All annotations in the final ground truth were manually verified and corrected and all networks trained for validation and benchmarking were trained de-novo from these final data. The mating and budding annotations of the 30 images in the test set were re-annotated by four different authors (DB, JM, FT and DH) of this study to allow us to compare the accuracy of YeastMate to inter-human variation (Supplementary Figure S6). The single cell masks were left as-is unless the re-annotator wanted to annotate a mating or budding event where not all cells are present in the segmentation mask (e.g., a small bud), in which case they manually added the missing object to the mask with a new label in napari.

#### CNN architecture and training

We use the Detectron2 (Wu *et al.*, 2019) implementation of Mask R-CNN (He *et al.*, 2017) with modifications to the mask head and mask loss to produce a multi-class segmentation (instead of the default binary segmentation) for each detected object.

In short, Mask R-CNN extends Faster R-CNN (Ren *et al.*, 2015), which performs object detection using a 2-step strategy: images are first processed by a *backbone* CNN to extract features, which are then fed into a region proposal network (RPN) that performs binary object/non-object classification for a predefined regular grid of bounding boxes, called *anchors*. The features of proposal boxes with the highest scores from the RPN are then resized to a common shape and fed into box classification and box regression *head* subnetwork, performing a second round of (multiclass) classification of the proposed regions, and refining the coordinates. Mask R-CNN adds a mask head to this architecture that produces a binary segmentation mask for each proposal region.

For YeastMate, we replaced the default mask head with a multiclass mask head: instead of a binary mask for each region, we produce a multichannel semantic segmentation (Supplementary Figure S1). We also do not threshold the segmentation, instead we continue working with the raw probabilities of the different segmentation classes $c_{seg}$. Note that the classes for the classification head $c_{obj}$, classifying the whole region, do not have to equal the classes of the segmentation. In our case, $c_{obj}$ classifies a whole region as single cell, mating or budding, whereas the segmentation mask has classes $c_{seg} \in$ {0: background, 1: single cell, 2: mating mother, 3: mating daughter, 4: budding mother and 5: budding daughter}.

During training, the single cell masks from the ground-truth (GT) data are used as one-hot encoded segmentations of class single cell and for all mating and budding events, we use masks encoding the role of all the individual cells (mother or daughter) during the event as GT. As our mask head performs multiclass classification, we use softmax activation in the last layer of the mask head and cross entropy (CE) loss instead of sigmoid activation and binary CE (BCE) loss.

We performed all training on a compute server equipped with 2 Intel(R) Xeon(R) E5-2680v4 CPUs (14 cores/28 threads each), 256GB of RAM and 2 NVIDIA Tesla V100 GPUs with 32GB VRAM each. We did not perform single training runs on multiple GPUs, instead using the two GPUs to process two cross-validation splits in parallel. For optimization of the network weights, we used stochastic gradient with momentum of 0.9 and a learning rate warm-up period of 1000 iterations. During cross-validation, the performance on the validation set stopped increasing after 200,000 iterations, we therefore also trained the final model using all training images for 200,000 iterations. We used a learning rate of 0.005 for the final model, based on performance during our cross-validation. Each training iteration was done on a batch of two random 400x400px image crops. Random augmentation was done each iteration in the form of random flips (50% chance for an upside-down flip and 50% for a left-right flip) and random rotation (a random 0-3 number of 90° rotations).

#### Postprocessing of CNN outputs

In a postprocessing step, we resolve the complementary segmentation and object detection outputs of our network to generate a single cell instance segmentation that assigns roles to cells participating a lifecycle transition event (Supplementary Figure S2).

For each detected bounding box of the compound classes mating or budding, we proceed as follows: we collect all detected objects of class single cell whose centers fall inside the compound object. We threshold the segmentation mask (for channel $c_{seg}$= single cell) of the single cell at 0.5 and calculate the average scores for all classes inside the mask in the segmentation of the compound object.

We consider the assignment of single cell objects to the available roles in a lifecycle transition object (I.e., mother and daughter in buddings, and two mothers and an optional daughter in a mating event) as a linear assignment problem and find the cells-to-roles matching with the highest overall score. For optional roles (e.g., the daughter in matings, as not all zygotes will have produced a medial bud), we discard the assignment of the cell to that role if the score lies under a threshold optional_object_thresh (we currently do not expose this parameter to the front end, instead using a value of 0.15, which was determined to be optimal during cross-validation). In each compound object, we also perform this matching for the other type and if the average score of the assignments is at least parent_override_thresh times higher, we use this assignment instead, thus using the segmentation output as a second line of evidence to catch zygotes erroneously classified as buddings by the object detection pathway, and vice-versa. During cross-validation we did not see strong improvements by using this overriding strategy, so we currently leave the parameter at a high default value of 20. All single cells that are not assigned a role in this procedure remain classified as single cells in the final segmentation.

Note that the procedure is written in a generalized way and could easily be adapted to accommodate, e.g., meiotic asci detection as well, if adequately annotated data were provided to our system.

#### Validation and Benchmarking of object detection performance

For validation of our models, we split our 147-image dataset into 5 parts of 29/30 images each, using a stratified splitting strategy placing a roughly equal number of images from individual experiments (Supplementary Table ST1) in each split. Four of the splits were used as cross-validation data to optimize network architecture and hyperparameters, while the final 30-image split was left aside as test data and only used in a final test run. Performance during cross-validation was assessed by training the network on 3 of the 4 training splits and using the remaining one as a validation set. The final model was trained on all 4 training splits and tested on the test split.

YeastMate provides bounding boxes of all detected objects as well as their scores. For each class of object, we sorted the detections by score (with no minimal score threshold) and generated precision-recall (PR) curves as well as average precision (AP) scores at 0.5 intersection-over-union (IoU) using the Python implementation of common object detection metrics in (Padilla *et al.*, 2021). We not only computed scores and PR-curves for the compound object classes mating and budding, but also for the roles of single cells within them (mating mother, mating daughter/medial bud, budding mother, budding daughter). For this, we assigned the single cell detection the score of the enclosing compound object before calculating performance metrics.

To assess reproducibility in mating and budding detection between different human annotators, we chose the first author (DB) as the reference annotator and compared the annotations of three other authors (JM, FT and DH). We assessed precision and recall for mating events, buddings, as well as the individual mother and daughter cells involved in them. For each class of object, the annotation from the second annotator was considered true positive (TP) if its bounding box overlapped an annotation by DB with an IoU of at least 0.5, otherwise, it was considered a false positive (FP). Annotations by DB that are not present in the re-annotation were considered false negatives (FN). For each class of object and each re-annotator, we calculated precision ($\frac{\#TP}{(\#TP + \#FP)}$) and recall ($\frac{\#TP}{(\#TP + \#FN)}$) and visualized them as points in the PR-curve describing the network performance (Supplementary Figure S6).

#### Single cell segmentation performance

To compare the single-cell segmentation performance of YeastMate to existing deep learning-based solutions, we applied our model (trained on our entire training dataset) as well as the pretrained models of YeastSpotter, YeaZ and YeastNet (Lu *et al.*, 2019; Dietler *et al.*, 2020; Salem *et al.*, 2021) to our test dataset, the brightfield images dataset provided by YeaZ as well as the dataset compiled by YeastNet, consisting of two novel annotated timeseries as well as segmentation masks for two of the timeseries provided by the Yeast Imaging Toolkit (YIT, http://yeast-image-toolkit.biosim.eu/pmwiki.php) .

Given a ground truth segmentation mask and a predicted segmentation mask, we matched detected cells to ground truth cells to maximize the total IoU of all GT-prediction pairs. We considered a predicted cell true positive (TP) if it had an IoU of over 0.5 with a ground truth cell and false positive (FP) otherwise. Ground truth cells that were not matched to a predicted cell with IoU > 0.5 were considered false negatives (FN). For each image, we calculated precision, recall and $F_1$-score ($\frac{2*(precision * recall)}{(precision + recall)}$) to measure object detection performance. In addition, we calculated the mean IoU of all true positive detections. If we tested multiple parameters for a tool, we only used the prediction with the highest $F_1$ score for each image.

##### YeastMate

We used YeastMate as a Python library to perform the benchmarking, but only varied parameters that are also available to end users via our graphical interfaces. For images with pixel sizes differing considerably from 110nm (our own images with 1.5x magnification and the images from YIT), we rescaled them using bilinear interpolation and then rescaled the mask produced by YeastMate to the original size using nearest-neighbor interpolation with no anti-aliasing. We normalized images based on their 0.015 and 0.985 quantiles before feeding them into our network. For each image, we generated segmentation masks with single cell score thresholds in the range (0.4, 0.5, …, 0.9).

##### YeastSpotter

We set up the standalone Python code of YeastSpotter (https://github.com/alexxijielu/yeast_segmentation) in a conda environment with the minimum required versions of tensorflow (1.10) and keras (2.2.4), as per the authors instructions. The only user-settable parameter in the main options file is a scale factor, but it can only be set to integer downsampling factors and is intended as a performance option to speed up inference. Therefore, we only used the default settings of no downsampling when making predictions with YeastSpotter.

##### YeaZ

We set up the standalone version of YeaZ (https://github.com/lpbsscientist/YeaZ-GUI) as per the authors instructions and followed the instructions for segmentation without using the GUI (using the provided weights for brightfield detection), omitting the tracking step. We applied YeaZ with different values for the two main parameters exposed in the GUI, threshold (0.3, 0.4, …, 0.9) and minimum distance between seeds (2, 5, 10).

##### YeastNet

We set up YeastNet (https://github.com/kaernlab/YeastNet) as per the authors instructions and used the main track.py script to process all datasets. No user-settable parameters are available via the command line interface, so we applied YeastNet with default parameters.

#### Application Architecture

YeastMate consists not only of the code for CNN-based cell detection and segmentation, but we also offer two separate graphical frontends for it to maximize user-friendliness: 1) standalone frontend written in React.js and packaged using Electron.js and 2) a plugin that can be used from the popular biomedical image analysis program Fiji (Schindelin *et al.*, 2012). An overview of the components of our software stack is shown in Supplementary Figure S3.

##### Detection Server and Python library

Our object detection and segmentation model is an extended version of the PyTorch Mask R-CNN implementation provided by Detectron2, combined with custom postprocessing routines to resolve single cells and lifecycle transitions. The detection model is wrapped via a

YeastMatePredictor class, which can load pretrained weights and network configurations and provides methods that will accept an image as input and return an integer-valued instance segmentation image as well as object detections in a custom JSON format. Upon installing the YeastMate module using pip, users can use functions from it in their own Python code. Note that the detection network expects intensities in the 0-1 range for the input images. To mitigate effects of individual bright or dark pixels, we normalize not based on minimal and maximal value, but rather a low and high, user-settable, intensity quantile (by default 0.015 and 0.985). Also, we found that for optimal performance, the input images should have a pixel size comparable to our reference pixel size of 110nm. We therefore also offer automatic resizing of images in the main inference method.

For using the YeastMate detector from other applications, such as our two GUI frontends, we wrap the detection pipeline as a webservice using Flask. Our detection server listens for multipart HTTP POST requests containing a 2-dimensional image (encoded as a single-channel 32-bit TIFF image) as well as parameters encoded as JSON (we follow the AnnotatedImageInput adapter API from BentoML, which we used for model serving initially, see https://docs.bentoml.org/en/latest/api/adapters.html#annotatedimageinput for details).

To ensure a reproducible environment with compatible versions of the libraries we use, we provide a Docker image of YeastMate, which can be used for running the detection server or re-training on new data.

#### React.js/Electron GUI client

To interface with our detection service, we created a custom GUI frontend based on modern web technologies (React.js) but provided as a desktop application using Electron.js. The frontend serves as a control hub to interface with other processes that perform image IO and visualization.

##### *GUI Frontend*

The frontend is written as a JavaScript web application using React.js and packaged as a desktop app using Electron.js (Supplementary Figure S4). It serves as a user interface for specifying all relevant settings of YeastMate and submitting an inference job to separate processes for image IO and the actual detection via HTTP requests. Users can launch the backend processes from the GUI or specify the IP addresses of already running instances (e.g. when the detection backend is running on a remote compute server)

##### *Client IO Backend*

The IO backend of our client is written in Python and is responsible for reading images from paths provided by the frontend (and in the case of multidimensional stacks, selecting the 2D plane the uses wishes to perform detection on), sending them to the detection server and handling and saving detection results. It is implemented as a webservice using Flask, with a task queue based on huey2. In addition to loading images and sending them to the detection server, it can also create crops of detected objects and save them to new files. It also provides several preprocessing functions that are helpful when dealing with multidimensional data.

##### *Preprocessing of images*

In the IO backend of our Electron client, we also support TIFF input images of higher dimensionality, given that they follow the TZCYX dimension order (the default order used by ImageJ/Fiji). The user can set a specific time point, z-plane, and channel, which will be sent to the detection server. Export operations such as cropping detected zygotes or budding events will be applied to all channels, timepoints and z-planes.

If data are provided in a different format, such as a microscope vendor-specific image format, we offer an optional preprocessing step that will resave the images as TIFF files meeting our requirements. Image IO from vendor specific formats is done via the BioFormats library (Linkert *et al.*, 2010) using the pims-bioformats wrapper. We tested this process extensively using Nikon nd2 files, but BioFormats should allow the reading of many other formats. As many of our own experiments were performed on a dual-camera microscope, during the resaving step we optionally added the option to align the color channels acquired on the two cameras if a reference channel (e.g., brightfield) was recorded on both. We perform an affine alignment using ORB keypoint detection and descriptor generation (Rublee *et al.*, 2011) followed by descriptor matching and outlier-resistant estimation of an affine transformation from matched keypoints using RANSAC (Fischler and Bolles, 1981) and will use the transformation to warp all channels from the second camera to match channels from the first. For these operations, we make use of the OpenCV and scikit-image packages. Note that for time series, we will align each frame individually and for z-stacks, we estimate a single 2D transformation on a maximum intensity projection of the reference stacks and applying the same transformation to all planes – we thus only correct misalignments along the xy-plane and will not correct drift in time series.

##### *Annotation Backend*

To visualize or correct the detections generated by the detection server, our GUI provides a visualization and annotation function. This will start a separate Python process that loads the specified input files, segmentation masks and JSON object annotations, which will be displayed using napari. Users can edit the masks and object annotations to manually curate the network output.

##### *All-in-one package*

We provide all-in-one packages for all major operating systems consisting of the Electron frontend as well as the IO backend and detection server, packaged using PyInstaller. When starting the Electron application and running a detection task, the IO backend and detection backend will be started in separate processes automatically (unless the user specifically points the frontend to the IP address and port of already running instances).

#### Fiji Plugin

In addition to our React-based frontend, we also provide a Fiji plugin that interfaces directly with a detection server using HTTP requests. The plugin is implemented as a SciJava Command (Rueden *et al.*, 2017) that acts on the currently selected image in Fiji (Supplementary Figure S5). It will transmit a normalized version of the image as well as detection parameters to the server and show the segmentation result as a new image and optionally add bounding boxes to Fijis ROI Manager. Execution of the plugin can be recorded as a macro for easy batch-application to many images. We implemented pixel-level calculations in the plugin using imglib2 (Pietzsch *et al.*, 2012).

#### User Guide

We provide a detailed user guide for YeastMate at https://yeastmate.readthedocs.io/

#### Code and data availability

YeastMate is fully open-source and licensed under the MIT license, all code is available on GitHub in the following repositories:

https://github.com/hoerlteam/YeastMate
(detection backend, **location of releases and issue tracking**)
https://github.com/hoerlteam/YeastMateFrontend (React.js/Electron.js GUI Application)
https://github.com/hoerlteam/YeastMateBackend
(Python-based IO and annotation code for the App)
https://github.com/hoerlteam/YeastMateFiji (Fiji plugin)

Pre-packaged binaries of the whole software stack are also available under:
https://github.com/hoerlteam/YeastMate/releases

Our final model weights and all training and testing data and annotations are available under:
https://osf.io/287fr/?view_only=99d1fddb563b4253957f226c19c4113f

### Supplementary Results

#### Single cell segmentation benchmark

When comparing YeastMate to other deep learning-based solutions for yeast cell instance segmentation, we achieve very satisfactory results (Supplementary Table ST2), outperforming all other tools on our own test dataset in all metrics considered. For the YeaZ and YeastNet-YIT datasets, we achieve the highest object detection performance (F1) and are only narrowly beat by YeaZ in terms of recall on the YeastNet-YIT dataset. In terms of the mean IoU of true positive detections, we achieve reasonable performance, but do not outperform YeaZ and YeastSpotter. Looking at example segmentation masks (Supplementary Figure S7), one can see a potential reason for our relatively low IoU: YeastMate tends to produce small segmentation masks compared to the other tools. So, while we capture the overall shape of cells well (except for very elongated mutants, which we likely mistake for zygotes consisting of two "mother cells", see Supplementary Figure S7E, an issue that could be remedied by including the YeaZ dataset into the training data), biases in the ground truth, e.g., whether to include the out-of-focus halo around cells into the mask, affect the performance of all tools. Overall, the Mask R-CNN-based tools YeastMate and YeastSpotter show a more consistent performance across datasets, a tribute to the robustness of the underlying network architecture (especially considering that YeastSpotter was not even trained on *S. cerevisiae* images), whereas U-Net-based (Ronneberger *et al.*, 2015) solutions seem to be more dependent on postprocessing steps optimized for a specific dataset. For example, YeastNet, which offers no easy way to finetune the watershed-based postprocessing, generalized poorly to data that do not match the imaging conditions of the original training dataset and seems prone to oversegmentation and artifacts, e.g., in the presence of uneven illumination.

### Author contributions

Conceptualization: DB, CO, DH

Code: DB, JM, DH

Experiments and image acquisition: FT, CJ

Annotation of images: DB, JM, FT, DH

DB and DH wrote the manuscript with input from all authors. All authors approved the final version.

### Supplementary References


Dietler,N. *et al.* (2020) A convolutional neural network segments yeast microscopy images with high accuracy. *Nat Commun*, **11**, 5723.

Fischler,M.A. and Bolles,R.C. (1981) Random sample consensus: a paradigm for model fitting with applications to image analysis and automated cartography. *Commun. ACM*, **24**, 381–395.

He,K. et al. (2017) Mask r-cnn. In, Proceedings of the IEEE international conference on computer vision., pp. 2961–2969.

Jakubke,C. *et al.* (2021) Cristae-dependent quality control of the mitochondrial genome. *Science Advances*, **in press**.

Linkert,M. *et al.* (2010) Metadata matters: access to image data in the real world. *The Journal of Cell Biology*, **189**, 777–782.

Lu,A.X. *et al.* (2019) YeastSpotter: accurate and parameter-free web segmentation for microscopy images of yeast cells. *Bioinformatics*, **35**, 4525–4527.

Padilla,R. *et al.* (2021) A comparative analysis of object detection metrics with a companion open-source toolkit. *Electronics*, **10**.

Pietzsch,T. *et al.* (2012) ImgLib2—generic image processing in Java. *Bioinformatics*, **28**, 3009–3011.

Ren,S. *et al.* (2015) Faster R-CNN: Towards real-time object detection with region proposal networks. In, Cortes,C. *et al.* (eds), *Advances in neural information processing systems 28.* Curran Associates, Inc., pp. 91–99.

Ronneberger,O. *et al.* (2015) U-Net: Convolutional Networks for Biomedical Image Segmentation. In, Navab,N. *et al.* (eds), *Medical Image Computing and Computer-Assisted Intervention – MICCAI 2015.* Springer International Publishing, Cham, pp. 234–241.

Rublee,E. *et al.* (2011) ORB: An efficient alternative to SIFT or SURF. In, *2011 International Conference on Computer Vision.* IEEE, Barcelona, Spain, pp. 2564–2571.

Rueden,C.T. *et al.* (2017) ImageJ2: ImageJ for the next generation of scientific image data. *BMC Bioinformatics*, **18**, 529.

Salem,D. *et al.* (2021) YeastNet: Deep-Learning-Enabled Accurate Segmentation of Budding Yeast Cells in Bright-Field Microscopy. *Applied Sciences*, **11**, 2692.

Schindelin,J. *et al.* (2012) Fiji: an open-source platform for biological-image analysis. *Nat Methods*, **9**, 676–682.

Sofroniew,N. *et al.* (2021) napari/napari: 0.4.11 Zenodo https://zenodo.org/record/5399494.

Wu,Y. et al. (2019) Detectron2. https://github.com/facebookresearch/detectron2.

### Supplementary Tables

Supplementary Table ST1: Dataset and Imaging Conditions

Supplementary Table ST2: Single cell segmentation performance

**Supplementary Table ST1:** Overview of images included in the YeastMate dataset. Unless noted otherwise, we took the middle z-plane from stacks acquired for (Jakubke *et al.*, 2021) to train and test our network.

| *Identifier* | *Cell types* | *Imaging conditions* | *#images* | *#single cells* | *#mating events* | *#budding events* | *Notes* |
|---|---|---|---|---|---|---|---|
| *wt_dic_1001/ wt_dic_1002* | WT ATP6-NG<br>x<br>WT matrix mKate2 | Ti2, 100x NA 1.49, DIC,<br>pixel size: 110nm | 40 | 5338 | 958 | 697 | |
| *dmic60_dic* | *Δmic60* ATP6-NG<br>x<br>*Δmic60* matrix mKate2 | Ti2, 100x NA 1.49, DIC,<br>pixel size: 110nm | 24 | 4125 | 735 | 928 | |
| *wt_bf_1/ wt_bf_2* | WT ATP6-NG<br>x<br>WT matrix mKate2 | Ti2, 100x NA 1.49, brightfield,<br>Low illumination intensity,<br>pixel size: 110nm | 40 | 4217 | 556 | 602 | Preliminary data for (Jakubke *et al.*, 2021) |
| *wt_bf_ magnified_rep1* | WT ATP6-NG matrix TagBFP<br>x<br>WT ATP6-mKate2 matrix TagBFP | Ti2, 100x NA 1.49<br>+ 1.5x magnification,<br>brightfield,<br>pixel size: 73nm | 5 | 339 | 8 | 123 | |
| *wt_bf_ magnified_rep2* | WT ATP6-NG matrix TagBFP<br>x<br>WT ATP6-mKate2 matrix TagBFP | Ti2, 100x NA 1.49<br>+ 1.5x magnification,<br>brightfield,<br>pixel size: 73nm | 4 | 248 | 5 | 100 | |
| *wt_bf_ magnified_rep3* | WT ATP6-NG matrix TagBFP<br>x<br>WT ATP6-mKate2 matrix TagBFP | Ti2, 100x NA 1.49<br>+ 1.5x magnification,<br>brightfield,<br>pixel size: 73nm | 19 | 983 | 64 | 384 | |
| *multiwell_bf_1* | WT ATP6-NG<br>x<br>WT matrix mKate2 | IX83, 60x NA 0.9, brightfield,<br>pixel size: 108nm | 9 | 872 | 52 | 330 | Preliminary data for high-throughput screen |
| *multiwell_bf_2* | WT ATP6-NG<br>x<br>WT matrix mKate2<br>/ *Δsrn2* matrix mKate2 | IX83, 60x NA 0.9, brightfield,<br>pixel size: 108nm | 6 | 936 | 2 | 451 | Preliminary data for high-throughput screen |
| *TOTAL* | | | 147 | 17058 | 2380 | 3615 | |

**Supplementary Table ST2:** Single cell detection (Precision, Recall, $F_1$) and segmentation (mean IoU of true positive detections) performance of YeastMate, YeastSpotter, YeaZ and YeastNet, with a 0.5 IoU threshold, shown as mean ± standard deviation across all images in a dataset. The best-performing tool per dataset and metric is highlighted by **bold** text. The results of YeaZ and YeastNet are grayed out for their respective datasets, as those images were used to train the models and thus the results do not reflect performance on novel data.

A: YeastMate test dataset

| | **PRECISION** | **RECALL** | **F1** | **MEAN IOU** |
|---|---|---|---|---|
| **YEASTNET** | 0.256 ± 0.279 | 0.220 ± 0.248 | 0.257 ± 0.236 | 0.557 ± 0.265 |
| **YEASTSPOTTER** | 0.775 ± 0.099 | 0.860 ± 0.070 | 0.812 ± 0.073 | 0.820 ± 0.030 |
| **YEAZ** | 0.670 ± 0.121 | 0.816 ± 0.072 | 0.733 ± 0.096 | 0.797 ± 0.026 |
| **YEASTMATE** | 0.945 ± 0.032 | 0.961 ± 0.031 | 0.953 ± 0.024 | 0.870 ± 0.019 |

B: YeaZ dataset

| | **PRECISION** | **RECALL** | **F1** | **MEAN IOU** |
|---|---|---|---|---|
| **YEASTNET** | 0.602 ± 0.318 | 0.609 ± 0.280 | 0.608 ± 0.281 | 0.733 ± 0.134 |
| **YEASTSPOTTER** | 0.965 ± 0.087 | 0.915 ± 0.114 | 0.938 ± 0.074 | 0.875 ± 0.058 |
| **YEAZ** | 0.991 ± 0.041 | 0.986 ± 0.045 | 0.988 ± 0.036 | 0.948 ± 0.025 |
| **YEASTMATE** | 0.967 ± 0.051 | 0.966 ± 0.044 | 0.966 ± 0.041 | 0.795 ± 0.044 |

C: YeastNet / YIT dataset

| | **PRECISION** | **RECALL** | **F1** | **MEAN IOU** |
|---|---|---|---|---|
| **YEASTNET** | 0.986 ± 0.039 | 0.890 ± 0.074 | 0.934 ± 0.047 | 0.856 ± 0.053 |
| **YEASTSPOTTER** | 0.724 ± 0.229 | 0.839 ± 0.113 | 0.749 ± 0.175 | 0.763 ± 0.075 |
| **YEAZ** | 0.681 ± 0.236 | 0.969 ± 0.053 | 0.771 ± 0.209 | 0.821 ± 0.064 |
| **YEASTMATE** | 0.918 ± 0.075 | 0.955 ± 0.051 | 0.935 ± 0.056 | 0.804 ± 0.076 |

### Supplementary Figures

Supplementary Figure S1: Network architecture

Supplementary Figure S2: Postprocessing of network output

Supplementary Figure S3: Software architecture

Supplementary Figure S4: Screenshots of Electron frontend

Supplementary Figure S5: Screenshots of Fiji frontend

Supplementary Figure S6: Object detection performance and comparison to human re-annotation

Supplementary Figure S7: Examples of single cell segmentation in comparison to existing tools

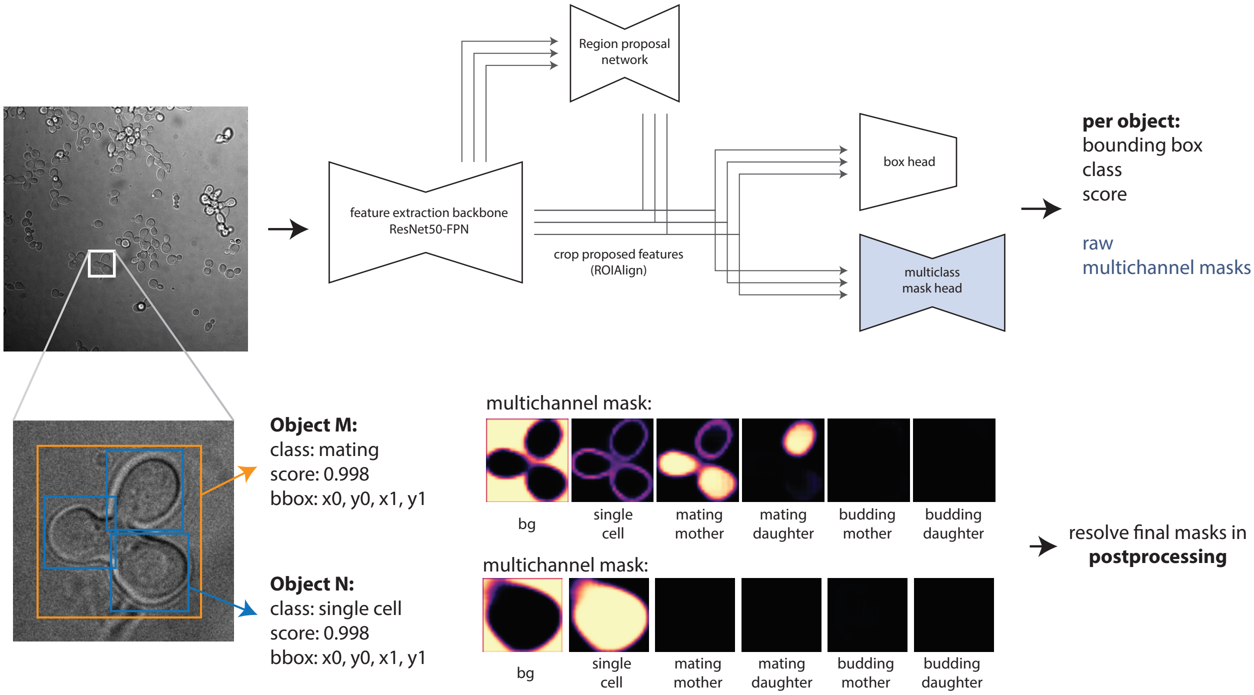


**Supplementary Figure S1:** Schematic Network architecture used in YeastMate: Images are fed through a modified Mask R-CNN with a ResNet50-FPN backbone (top). In addition to object detections with bounding boxes with object classes $c_{obj} \in$ {single cell, mating, budding}, our modified mask segmentation head produces multichannel masks with segmentation classes $c_{seg} \in$ {background, single cell, mating mother, mating daughter, budding mother, budding daughter} (bottom).

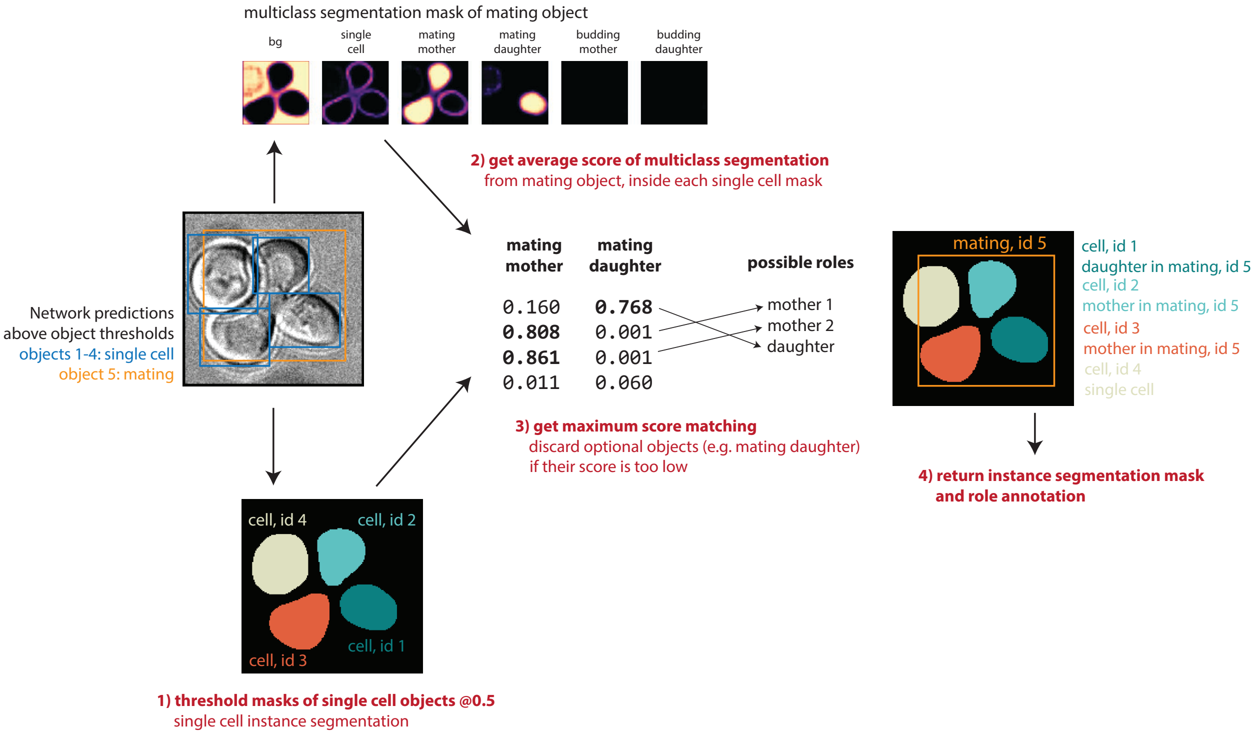


**Supplementary Figure S2:** YeastMate Postprocessing pipeline: We take the outputs of our modified Mask R-CNN (objects with bounding boxes, scores and classes $\in \{single\ cell, mating, budding\}$, as well as raw segmentation masks with classes $\in \{background, single\ cell, mating\ mother, mating\ daughter, budding\ mother, budding\ daughter\}$ ) and perform the following steps: 1) for all single cell objects (with score above a user-settable threshold) we threshold the mask of the "single cell" class at 0.5, which already gives us an instance segmentation of individual cells 2) for objects of classes mating or budding, we then get the mean score for respective subclasses within the single cell mask of each cell in the bounding box and 3) calculate an optimal assignment to the "roles" in a lifecycle transition event, which gives us 4) the final results consisting of an instance segmentation with additional annotations for transition events.

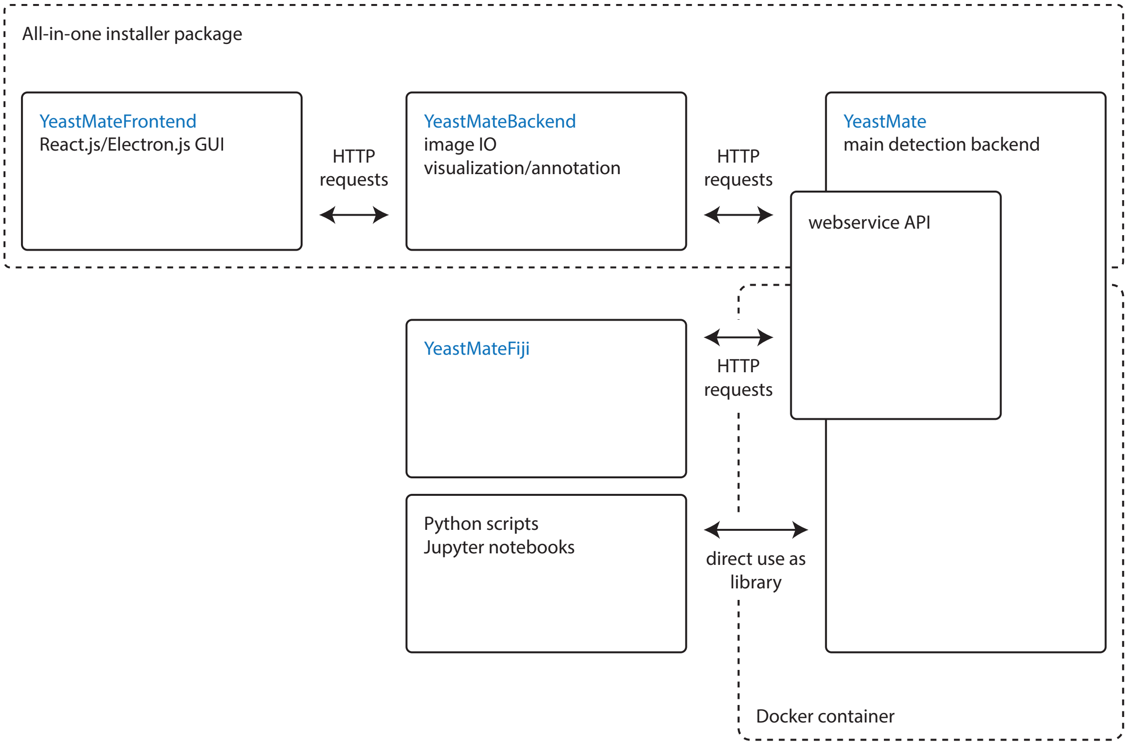


**Supplementary Figure S3:** Software components of YeastMate (blue components indicate individual repositories available on GitHub): At its core lies the detection backend (YeastMate, right), which contains our modified Mask R-CNN implemented using PyTorch and Detectron2. It can either be installed as a Python package and directly used from custom scripts (examples are provided in the repository). The detection backend can also be served as a webservice, which provides an interface for our two GUI frontends: the Fiji plugin YeastMateFiji (middle) as well as our custom GUI frontend (top). This frontend consists of the actual UI written in React.js and packaged as a desktop application using Electron.js (YeastMateFrontend) which will spawn Python-based subprocesses for Image IO, visualization using napari or communication with the detection server. This functionality is implemented in the YeastMateBackend repository. For convenience, we compile YeastMateFrontend, YeastMateBackend and YeastMate into a single installer package. We also offer a Docker image for the YeastMate package for re-training networks or serving the detection backend from a reproducible environment.

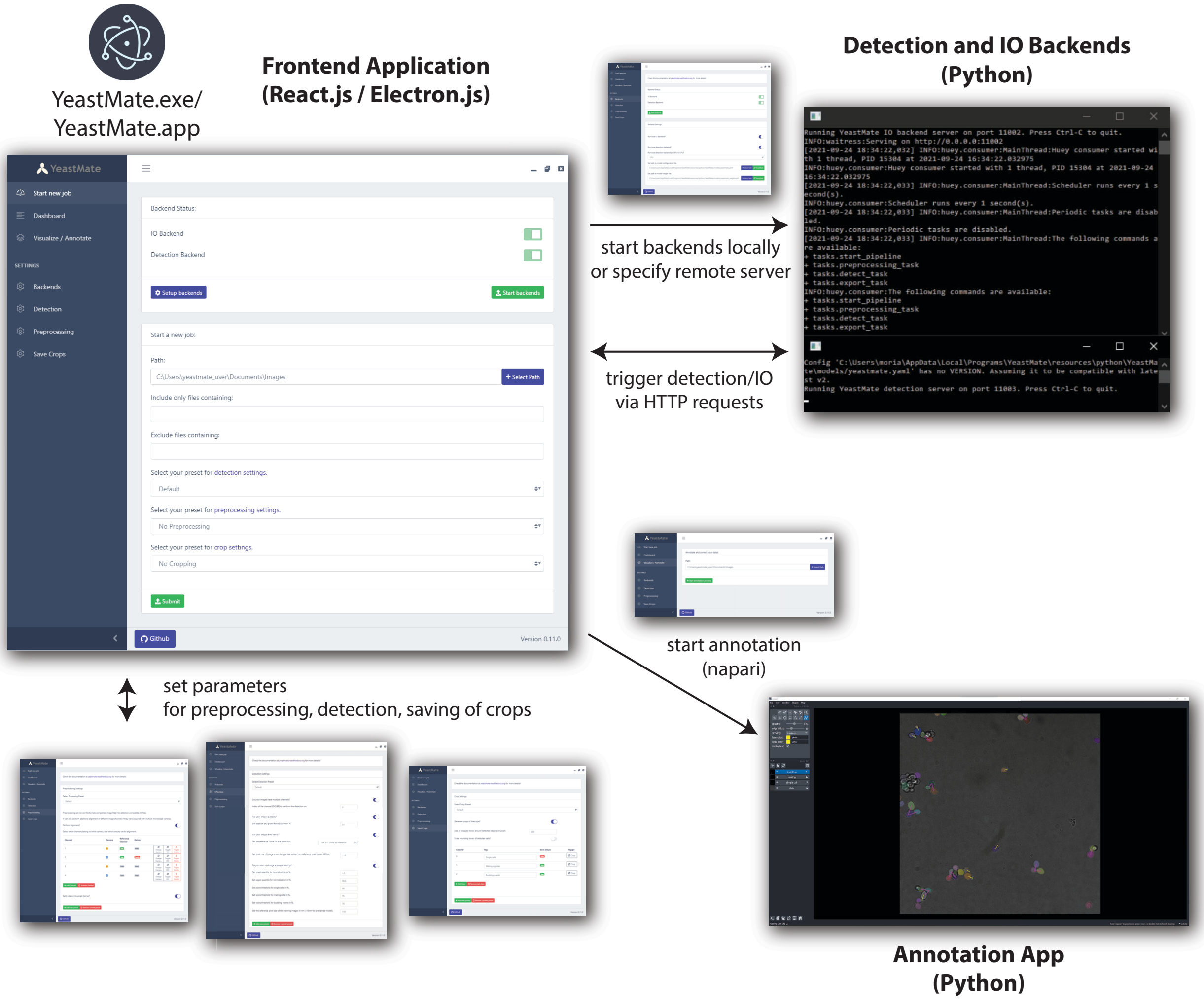


**Supplementary Figure S4:** YeastMate standalone application. We package all components of YeastMate in an all-in-one package consisting of a GUI frontend based on React.js and Electron.js. The GUI forms an interface to separate Python-based processes for image IO (including preprocessing of non-TIFF files to meet our format requirements and export of crops around detected objects), detection and annotation. The respective backend processes will be started automatically by the frontend.

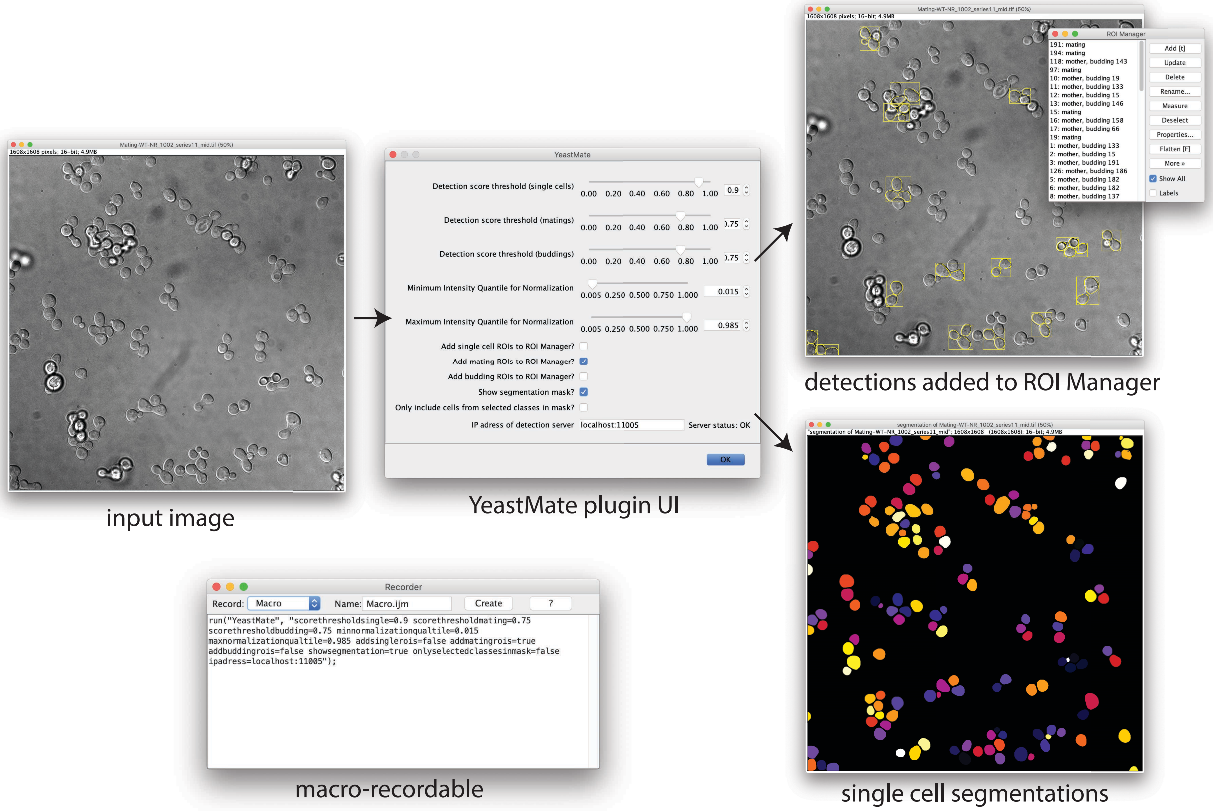


**Supplementary Figure S5:** Fiji-plugin frontend to YeastMate. The plugin can be from the Plugins menu under Plugins->YeastMate. Its GUI consists of a single dialog allowing the user to set score thresholds, normalization quantiles and what kind of output they want to display, as well as the IP of the detection server (a status message in the dialog will indicate if the server is reachable). Upon clicking "OK", the plugin will connect to the detection server and transmit the currently active image for detection. Once it receives the results, it will add the bounding boxes or outlines of the detected objects (or just objects of specific classes, if desired by the user) to Fijis ROI Manager and display a single cell segmentation mask. If the corresponding options are set, the mask can be limited to cells participating in mating or budding events. The whole process can be recorded using Fijis Macro Recorder and included in macros for, e.g., batch processing of multiple files.

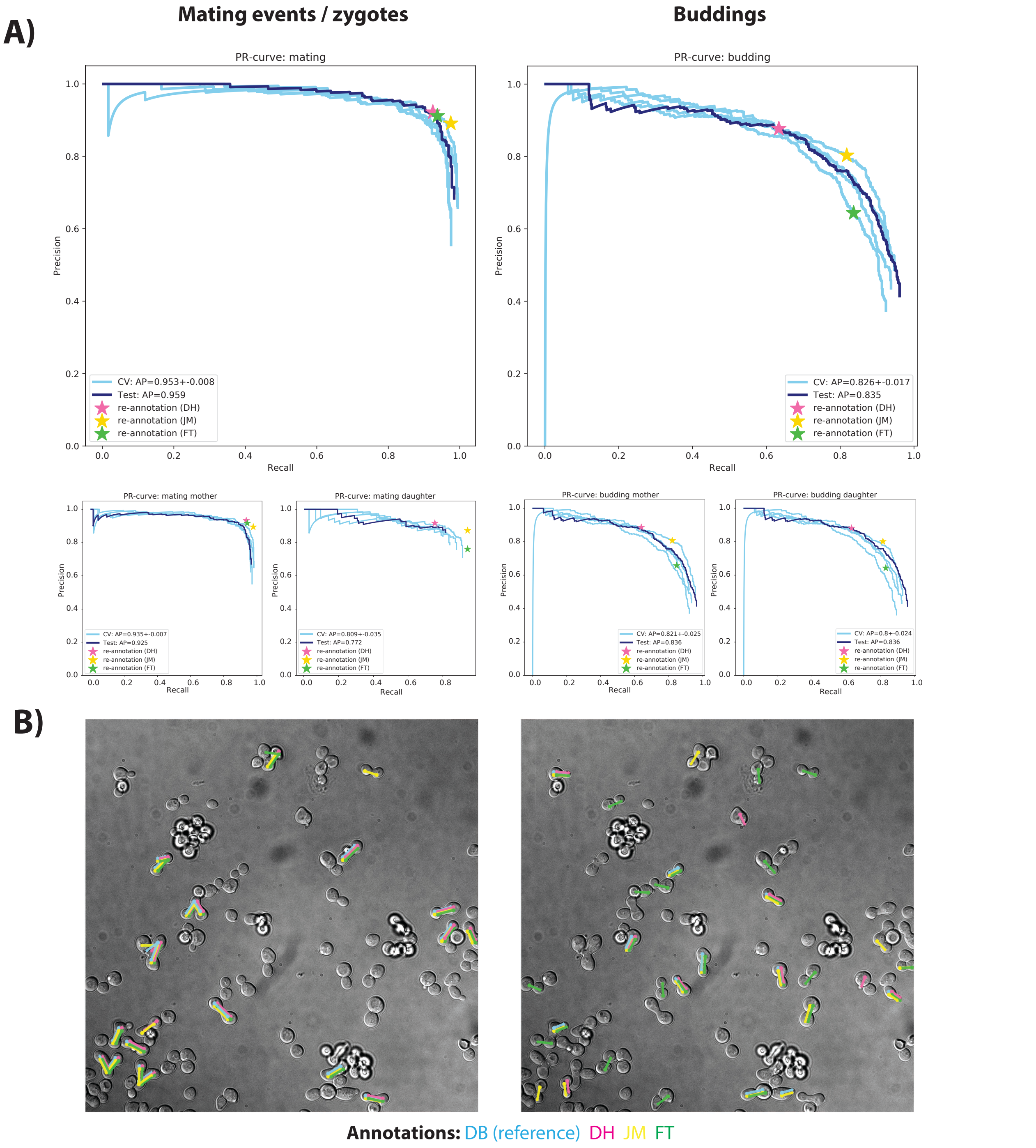


**Supplementary Figure S6:** Object detection performance. A, top) Precision-Recall curves for object detection performance (at 0.5 IoU threshold) for mating and budding events for the 4 cross-validation splits (light blue) and the final inference on the held-out test dataset (dark blue). Human re-annotation precision and recall for 3 re-annotators are shown as colored stars and closely coincide with the network curves. A, middle) Same curves as above, but for the mother and daughter subclasses of mating events and buddings. B): Visualization of the mating and budding annotations of the 4 annotators on an example image from the test set. Mating events are indicated by lines between the mother cells forming the zygote (circles) and, optionally, to a medial daughter bud (star). Buddings are indicated by a line from mother (circle) to daughter (star) cells. Note that shifts between the annotations of the different authors, as well as a random z-ordering of the annotations were introduced on purpose for this figure to make overlapping annotations easier to see.

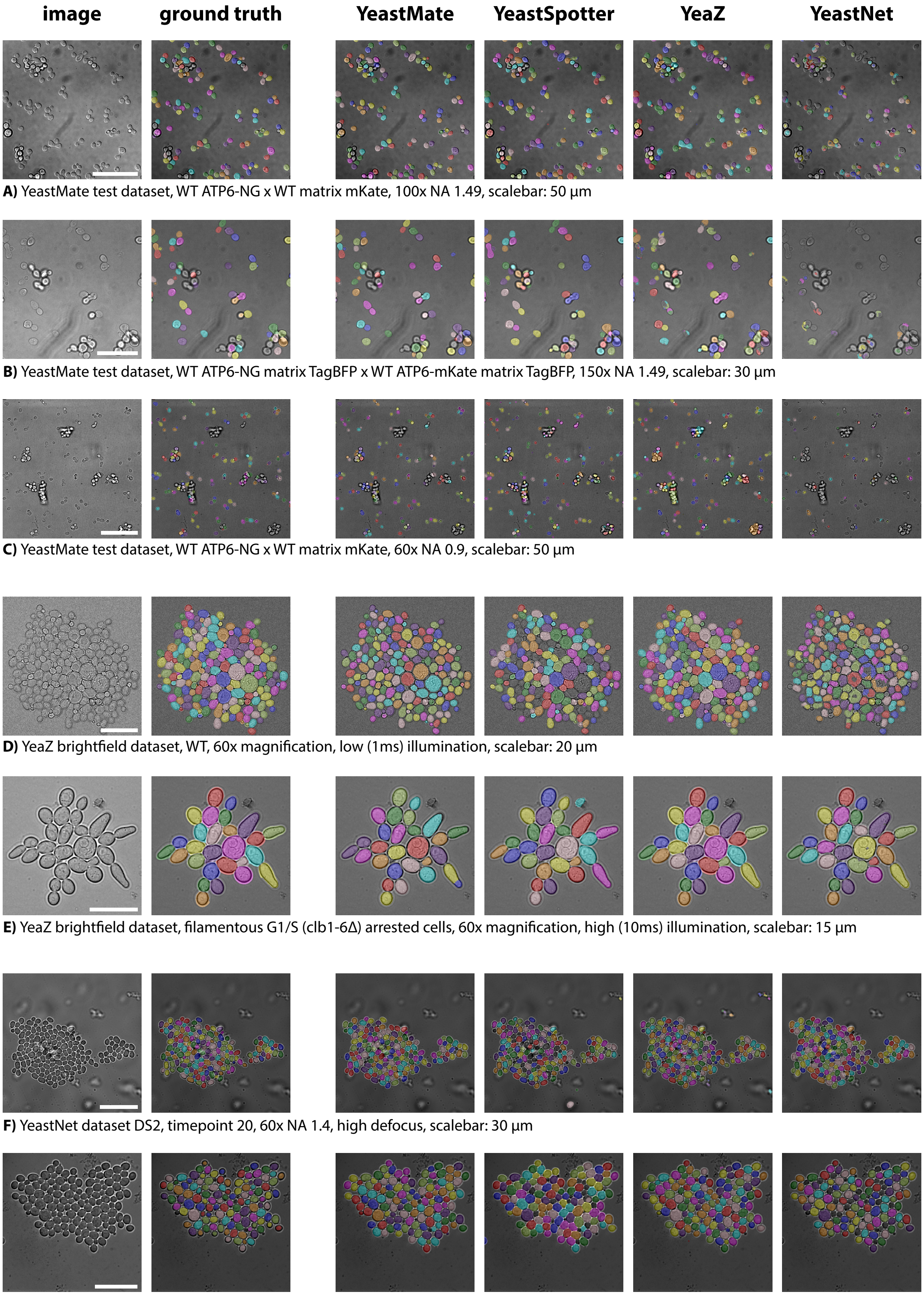


**A)** YeastMate test dataset, WT ATP6-NG x WT matrix mKate, 100x NA 1.49, scalebar: 50 µm

**B)** YeastMate test dataset, WT ATP6-NG matrix TagBFP x WT ATP6-mKate matrix TagBFP, 150x NA 1.49, scalebar: 30 µm

**C)** YeastMate test dataset, WT ATP6-NG x WT matrix mKate, 60x NA 0.9, scalebar: 50 µm

**D)** YeaZ brightfield dataset, WT, 60x magnification, low (1ms) illumination, scalebar: 20 µm

**E)** YeaZ brightfield dataset, filamentous G1/S (clb1-6Δ) arrested cells, 60x magnification, high (10ms) illumination, scalebar: 15 µm

**F)** YeastNet dataset DS2, timepoint 20, 60x NA 1.4, high defocus, scalebar: 30 µm

**G)** YIT dataset 3, timepoint 20, 100x NA 1.4, scalebar: 25 µm

**Supplementary Figure S7:** Single cell segmentation performance and comparison to existing deep learning-based tools for yeast segmentation on example images from our own test dataset (A-C) the YeaZ brightfield dataset (D,E) as well as the YeastNet/YIT dataset (F,G). Masks are overlaid on the original images in random colors using the label2rgb function from scikit-image.
